# Retinal affectation in Huntington’s disease mouse models concurs with a local innate immune response

**DOI:** 10.1101/2023.01.17.523578

**Authors:** Ana I. Arroba, Andrea Gallardo-Orihuela, Fátima Cano-Cano, Francisco Martín-Loro, María del Carmen González-Montelongo, Irati Hervás-Corpión, Pedro de la Villa, Lucía Ramón-Marco, Laura Gómez-Jaramillo, Luis M. Valor

**Author notes:** **Corresponding coauthors** Ana I. Arroba, Unidad de Investigación, 9th floor Hospital Universitario Puerta del Mar, INiBICA, Av. Ana de Viya 21, 11009 Cádiz, Spain., Luis M. Valor, Unidad de Investigación, 6th floor Diagnostics Building, Hospital General Universitario Dr. Balmis, ISABIAL, Av. Pintor Baeza 12, 03010 Alicante, Spain., Telf.

## Abstract

Huntington’s disease (HD) is a devastating disorder caused by aberrant expansion of CAG repeats in the *HTT* gene. Striatal dysfunction has been widely studied in HD mouse models. However, cumulative evidence indicates that the retina can also be functionally altered with consequences for visual function and circadian rhythms. The retina is the most exposed part of the central nervous system that can be used for monitoring the health status of patients using noninvasive techniques. To establish the retina as an appropriate tissue for HD studies, we linked the retinal alterations with those in the inner brain. We confirmed the malfunction of the R6/1 retinas, which underwent a rearrangement of their transcriptome as extensive as in the striatum, indicating a profound retinal affectation in HD. Tissue-enriched genes were downregulated in both areas, but a neuroinflammation signature was specifically induced in the R6/1 retina due to glial activation that was reminiscent of the situation in HD patient’s brains. These phenomena were confirmed in the zQ175 strain, and were accompanied by a differential impairment of the autophagy system between both tissues. Overall, these results demonstrated the suitability of the mouse retina as a research model for HD.

## Introduction

Huntington’s disease (HD) (OMIM #143100) is a fatal rare disorder without a cure, and its estimated prevalence is 1-15 per 100,000 people worldwide ^1^. HD is caused by an aberrant expansion (>39) of the polymorphic tract of CAG repeats at exon 1 of the *HTT* locus, which triggers a progressive symptomatology, usually starting in mid-adulthood (35-45 years old) that encompasses cognitive and motor impairments, psychiatric disorders and other disturbances until final death ^2^. Although the most prominent signs are derived from malfunctioning and degeneration of the basal ganglia and the corticostriatal circuitry ^3^, the involvement of other brain areas and peripheral tissues/cells plays a substantial role in the decline of quality of life, extending the repertoire of HD symptoms ^4–8^. However, our current toolbox of biomarkers to explain the large variability in the pleiotropic manifestation of symptoms with potential application in clinics is still limited ^9–11^, and justifies the search for novel biomarkers with prognostic value in HD to enable the evaluation of therapeutic responses and to facilitate decision-making during clinical management.

The retina is a highly organized tissue of the CNS characterized by high cellular diversity arranged in discrete layers. This organization determines the functionality of the retina, in which photoreceptor neurons (rods and cones) transduce light stimuli into the complex network of interneurons (amacrine, bipolar and horizontal cells) to finally converge into ganglion cells, which axons form the optic nerve ^12^; glial cells (astrocytes, Müller glia and resident microglia) and other nonneural types provide homeostatic and regulatory support to this network ^13^. The accessibility of the retina to noninvasive techniques offers unique opportunities to infer the health status of inner regions of the CNS in neurodegenerative disorders, assuming that the retina can reproduce to some extent the disruption of key mechanisms that compromise cell function and viability. In patients, the suitability of retinal imaging in diagnosis is being investigated in Alzheimer’s disease, Parkinson’s disease, Lewy body dementia, frontotemporal dementia and multiple sclerosis, with promising prospects ^14,15^. HD is not an exception; there are selective abnormalities (e.g., thickness reduction of the temporal retinal nerve fibre layer (RNFL), colour vision impairment) that can be tentatively correlated with performance on the Unified Huntington’s Disease Rating Scale (UHDRS). These anomalies can be accompanied by an additional affectation of the visual pathway, i.e., reduced amplitude of visual evoked potentials and impaired temporal contrast sensitivity (reviewed in ^16^). In the case of HD mouse models, mutant retinas express mHTT, display deeply reduced photoexcitability responses and undergo a cellular remodelling ^17–21^ that coincides with early stages of motor impairment: therefore, retinal affectation could be correlated with striatal impairment in HD.

However, the molecular mechanisms that compromise the retinal structure and visual function in HD mice and their corresponding correlates with inner brain alterations have not yet been investigated. This study reports for the first time a comprehensive analysis of the concomitant molecular alterations in the retina and striatum of HD mouse models, the transgenic R6/1 and the knock-in (KI) zQ175 strains, in which glial activation becomes a distinctive feature between both tissues during pathology progression.

## Materials and methods

### Animals

Transgenic R6/1 ^22^, KI zQ175 ^23^ and *rd10* mice ^24^, together with control wild-type (wt) animals, were maintained on a pure C57BL/6J background under a 12-h light/dark cycle with food and water provided *ad libitum*. The *rd10* mouse model of retinal degeneration was homozygous for the Pde6brd10 mutation and was kindly provided by Bo Chang (the Jackson Laboratory). Experimental protocols were approved by the Comité de Ética de Experimentación Animal - Órgano Habilitado de la Universidad de Cádiz and authorized by the Dirección General de la Producción Agrícola y Ganadera de la Junta de Andalucía according to European and regional laws.

### Spectral-domain optical coherence tomography (SD-OCT)

R6/1 (25-week-old), zQ175 (12-month-old) and matched-age wt mice were anaesthetized with ketamine (95 mg/kg) and xylazine (5 mg/kg) and maintained on a heated pad at 37°C. The eyes were instilled with a topical drop of 1% tropicamide (Colircusí Tropicamida, Alcon Cusí SA, Barcelona, Spain) for pupil dilation and 2% Methocel (Ciba Vision AG, Hetlingen, Switzerland). OCT images were obtained using a Micron IV rodent imaging system (Phoenix Research Labs, Pleasanton, CA, USA) as described elsewhere ^25^. A B-scan including the maximal retinal thickness (in the centre of the retina) was segmented between the inner limiting membrane and the base of the retinal pigment epithelium using the Insight software package (Phoenix Research Labs). The thickness of both eyes (containing average raw data from 624 A-scans from each eye after removing the first and last 200 scans) was averaged per animal for statistical analysis.

### Electroretinogram recordings

All electroretinogram (ERG) procedures have been described previously ^26^. Dark-adapted mice were anesthetized under dim red light and the eyes were instilled as in the SD-OCT procedure. All recordings were performed with the animal placed in a Faraday cage. Flash-induced ERG responses were recorded from the right eye in response to light stimuli produced with a Ganzfeld stimulator. Light intensity was measured in the eye with a photometer (Mavo Monitor USB, Nuremberg, Germany). A total of 4 to 64 consecutive light stimuli were presented. The interval between light flashes applied in scotopic conditions was 10 seconds for dim flashes and up to 60 seconds for the highest intensity; under photopic conditions, the interval between light flashes was fixed at 1 second. The ERG signals were amplified and band filtered between 0.3 and 10.000 Hz with a Grass amplifier (CP511 ACamplifier; Grass Instruments, Quincy, MA, USA). Electrical signals were digitized at 10 kHz with a Power-Lab data acquisition board (ADInstruments, Chalgrove, UK). Bipolar recordings were obtained using corneal electrodes (Burian-Allen electrode, Hansen Ophthalmic Development Laboratory, Coralville, IA, USA) and a reference electrode located in the mouth, whereas the ground electrode was located in the tail. Rod-mediated responses were recorded under dark adaptation; light flashes ranging from −4 to 1.5 log Cd × s × m^−2^ were used. Light flashes ranging from −1.5 to 0.5 log Cd × s × m^−2^ were used for recording of the mixed rod- and cone-mediated responses. Cone function was tested in response to light flashes ranging from −0.5 to 2 log Cd × s × m^−2^on a rod saturating background of 30 Cd × s × m^−2^. Measurements of wave amplitudes were estimated blinded to the experimental genotype of the animal.

### Nucleic acid extraction and PCR assays

Animals were sacrificed by cervical dislocation, and the striatum and retina were immediately dissected and submerged in RNAlater (ThermoFisher, Madrid, Spain) until processing. Total RNA and genomic DNA were sequentially extracted using TRIzol (Thermo Fisher, Madrid, Spain), and we next followed the procedures for RT-qPCR and CAG repeat analysis described in ^27^ with minor modifications: qPCR was performed on the Rotor-Gene 6000 Detection System (Corbett, Hilden, Germany) and QuantStudio 12K Flex (Thermo Fisher, Madrid, Spain), and each independent reaction was normalized to the level of *Tbp*, since its expression in the retinal samples was less variable across time points (coefficient of variation CV = 0.51) than other housekeeping genes (e.g., *Gapdh*, CV= 0.69; *Actb*, CV = 1.00); in addition, the latter genes were found to be significantly altered in the RNA-seq analysis (see below). Fold changes were estimated using the ΔΔC_T_ method. The sequences of all primer pairs are provided in Table S1.

### Immunohistochemistry assays

Eye balls and whole retinal explants were fixed in 4% paraformaldehyde in 0.1 M phosphate buffer, pH 7.4, at 4°C overnight and subsequently put in a solution of PBS with 25% sucrose for another 24h and cryopreserved in Tissue-Tek (4583, Sakura Finetek, Barcelona, Spain) at −80°C until cryosectioning. For immunofluorescence analysis, cryosections were incubated with a permeabilization solution containing 0.1 M TBS, 2% Triton X-100, blocked for 2 hours in TBS containing 3% BSA and 1% Triton X-100, and incubated overnight in a humid chamber at 4°C with rabbit anti-glial fibrillary acidic protein (GFAP) antibody (1:500, Z0334, DAKO, Agilent, Glostrup, Denmark), rabbit anti-ionized calcium binding adaptor molecule 1 (Iba-1) (1:500, 019-19741, WAKO, FUJIFIL Cellular Dynamics, Madison, WI, USA) or rabbit anti-huntingtin D7F7 (1:300, Cell Signaling Technology, Danvers, CA, USA) in blocking solution. Next, sections were washed and incubated for 2 h with secondary antibodies conjugated to Alexa-488 (1:2000; Molecular Probes, Thermo Fisher Scientific, Waltham, MA, USA). For the D7F7 antibody, the incubation was conducted with anti-rabbit biotin (1:300, B8895, Sigma-Aldrich, Darmstadt, Germany) followed by incubation with streptavidin-Alexa 488 (1:600, S11223, Thermo Fisher, Madrid, Spain), 2 hours each. After washing, both the retinal explants and sections were mounted with medium (Fluoromount G, Southern Biotech, Thermo Fisher Scientific, Waltham, MA, USA) containing 4’-6-diamidino-2-phenylindole (DAPI). Staining was observed with an inverted laser confocal microscope Axio Observer LSM900 (Carl Zeiss Microscopy GmbH, Göttingen, Germany). Cell nuclei labelled with DAPI were counted in the ONL using ImageJ software on retinal section images at 40× magnification. In each mouse, we measured a total of six retinal cross-sections made through the optic nerve head, and averaged for each animal.

### Cell death detection

Programmed cell death was determined in retinal cryosections by terminal deoxynucleotidyl transferase-mediated dUTP nick-end labelling (TUNEL) using the DeadEnd Fluorometric TUNEL system following the manufacturer’s instructions (Promega, Madison, WI, USA). Next, the retinas were mounted in Fluoromount G containing DAPI and observed with an inverted laser confocal microscope Axio Observer LSM900 (Carl Zeiss Microscopy GmbH, Göttingen, Germany). Serial optical sections were acquired with a 10×objective every 5 μm in depth around the optic nerve head. To ensure that TUNEL-positive nuclei were located in the ONL, the depth of analysis was set according to ONL thickness, as determined in retinal sections.

### Western blotting assays

Whole retinas were homogenized in lysis buffer containing 125 mM Tris-HCl pH 6.9, 2% SDS, and 1 mM DTT supplemented with protease inhibitors (cOmplete EDTA-free, Sigma-Aldrich, Darmstadt, Germany). All debris was removed by centrifugation at 14,000×g for 10 min at 4 °C and the protein concentration was quantified using the Bio-Rad protein assay with BSA as a standard. Equivalent amounts of protein were resolved using denaturing sodium dodecyl sulphate-polyacrylamide gel electrophoresis (SDS-PAGE), followed by transfer to PVDF membranes (Merck Millipore, Cork, Ireland). Membranes were blocked using 5% nonfat dried milk or 3% BSA in 10 mM Tris-HCl pH 7.5, 150 mM NaCl, and 0.1% Tween-20, and incubated overnight with several primary antibodies (1:1000) in fresh blocking solution. After incubation with secondary antibodies (1:5000), immunoreactive bands were visualized using enhanced chemiluminescence reagent (Bio-Rad, Hercules, CA, USA). The fold change relative to the basal condition is shown. Blots were quantified by scanning densitometry. Primary antibody against LC3 (#2775) were purchased from Cell Signaling Technology (Danvers, CA, USA); primary antibody against α-tubulin (T5168) and secondary antibodies (A0545, anti-rabbit; A9044, anti-mouse IgG-peroxidase) were purchased from Sigma-Aldrich (Darmstadt, Germany).

### RNA-seq analysis, external datasets and bioinformatics

After the TRIzol procedure, striatal and retinal RNA from the same R6/1 mice was pooled (3-4 samples per genotype) and further processed using the clean-up protocol of the RNeasy Mini Kit (Qiagen, Hilden, Germany), which also included on-column DNase I treatment. DNA libraries were produced for mRNA using the TruSeq Stranded mRNA kit (Illumina, San Diego, CA, USA) and subsequently sequenced using a NovaSeq apparatus (Illumina, San Diego, CA, USA) at STAB-VIDA facilities in a 150-bp paired-end configuration. The resulting reads (>40 M / sample) were mapped onto the mouse genome GRCm38/mm10 using “Salmon” software ^28^. Genes with <10 counts in all samples were removed. The normalization of read counts and differential expression analysis was conducted using the “DEseq2” package ^29^. Differentially expressed genes (DEGs) were filtered with an FDR threshold (adjusted *p-*value) of 0.05. The RNA-seq data can be downloaded from the Gene Expression Omnibus (GEO) database using the accession number GSE216520.

To determine the most affected cell subtypes by the expression of mHTTin the differential expression between R6/1 and wild-type littermates, we used the cell-specific signatures obtained from mouse retinas in physiological conditions ^30^, i.e., the genes that define the different subpopulations (clusters) of cells identified by scRNA-seq (*Supplementary Table S4* of the original publication ^30^). Amacrine and bipolar cells were defined by several clusters that we pooled for our study. Because rod cells constitute ~70% of the total retinal cells, contamination from rod cytoplasmic mRNA was present in nearly all clusters ^31^; therefore, we filtered out the rod signature of the cell-specific signatures before their use in our gene expression profiles. To detect potential differences in the cell content between genotypes, we used curated lists of 100-gene markers for neurons, astrocytes, endothelial cells and microglia that are shared between humans and mice and that have been further validated in the literature (*Supplemental File 2* of the original publication ^32^). To investigate the presence of glial activation markers, we used the most significant DEG (top 250): (i) between control and activated Aldh1h^+^-astrocytes (isolated from 1 to 7 days after different *in vivo* injuring protocols ^33^), as calculated by applying the “affy” ^34^ and “limma” ^35^ packages, and (ii) among CD11b^+^-CD45^+^ microglial cells isolated from the neurodegenerative model CK-p25 at different time points after induction of the transgene (from immediately to 6 weeks), as presented in *Supplementary Table S4* of the original publication ^36^. Astrocytic panmarkers and specific markers for neurotoxic (A1) and neuroprotective (A2) astrocytes were obtained from a previous report ^37^. We also used two classifications of autophagy-related genes as compiled by a recent study ^38^: one consisted of “mTOR and upstream pathways”, “autophagy core”, “autophagy regulators”, “mitophagy”, “docking and fusion”, “lysosome” and “lysosome-related”, and the other consisted of “autophagy induction” and “lysosomal biogenesis” genes. Overlapping genes between our RNA-seq analysis and external datasets are found in Supplementary Table S3.

Other additional bioinformatic tools included Venny (http://bioinfogp.cnb.csic.es/tools/venny/) for the identification of overlapping genes between multiple lists of genes, DAVID 2021 Update ^39^ for overrepresentation analysis of Gene Ontology (GO) terms related to biological processes, Pscan ^40^ for overrepresentation analysis of transcription factor binding sites (TFBS) at the promoter regions (−950/+50) from the Jaspar 2020_NR database, and the native R environment for statistical analysis (ANOVA and Student’s *t*-test).

## Results

### R6/1 retinas were morphologically and functionally altered

To confirm that our R6/1 colony also exhibited the retinal impairments reported in HD mouse models ^17–21^, we first demonstrated that mHTT was effectively expressed in the retina (Supplementary Fig. S1A). Using SD-OCT, 25-week-old R6/1 mice showed aberrant fundus images due to the presence of bright white or hyperreflective spots, as already reported ^20^, which might be indicative of inflammatory processes ^41–43^ (Fig. 1A). Similar aberrant fundus was observed in 12-month-old zQ175 retinas (Fig. 1B). This finding was accompanied by a reduction in the total retinal thickness in R6/1 mice, which was especially remarkable at the ONL since this layer also showed a significant thinning in zQ175 retinas (Fig. 1C) and a decrease in the number of nuclei compared to matched-age controls in both strains (Fig. 1D). However, this cell reduction was not linked with an increase in TUNEL staining (Supplementary Fig. S1B), which might be explained by the fact that cell death in HD brains does not necessarily follow the conventional apoptotic pathway of blebbing and fragmentation of the nucleus and cytoplasm ^44^. We also evaluated the visual function by ERG recordings of scotopic (dark-adapted) and photopic (light-adapted) responses to light flashes that are prominently mediated by rod and cone pathways, respectively. Symptomatic 24-week-old R6/1 mice showed a nearly complete abolishment in the amplitude of the a- and b-waves in both scotopic and photopic ERG, indicating affectation of both types of photoreceptor cells (a-wave) and downstream postsynaptic transmission (b-wave) (Fig. 1E). Overall, our results indicated that the retinas of R6/1 and zQ175 mice presented profound morphological alterations associated with visual response failure.

**Figure 1.**
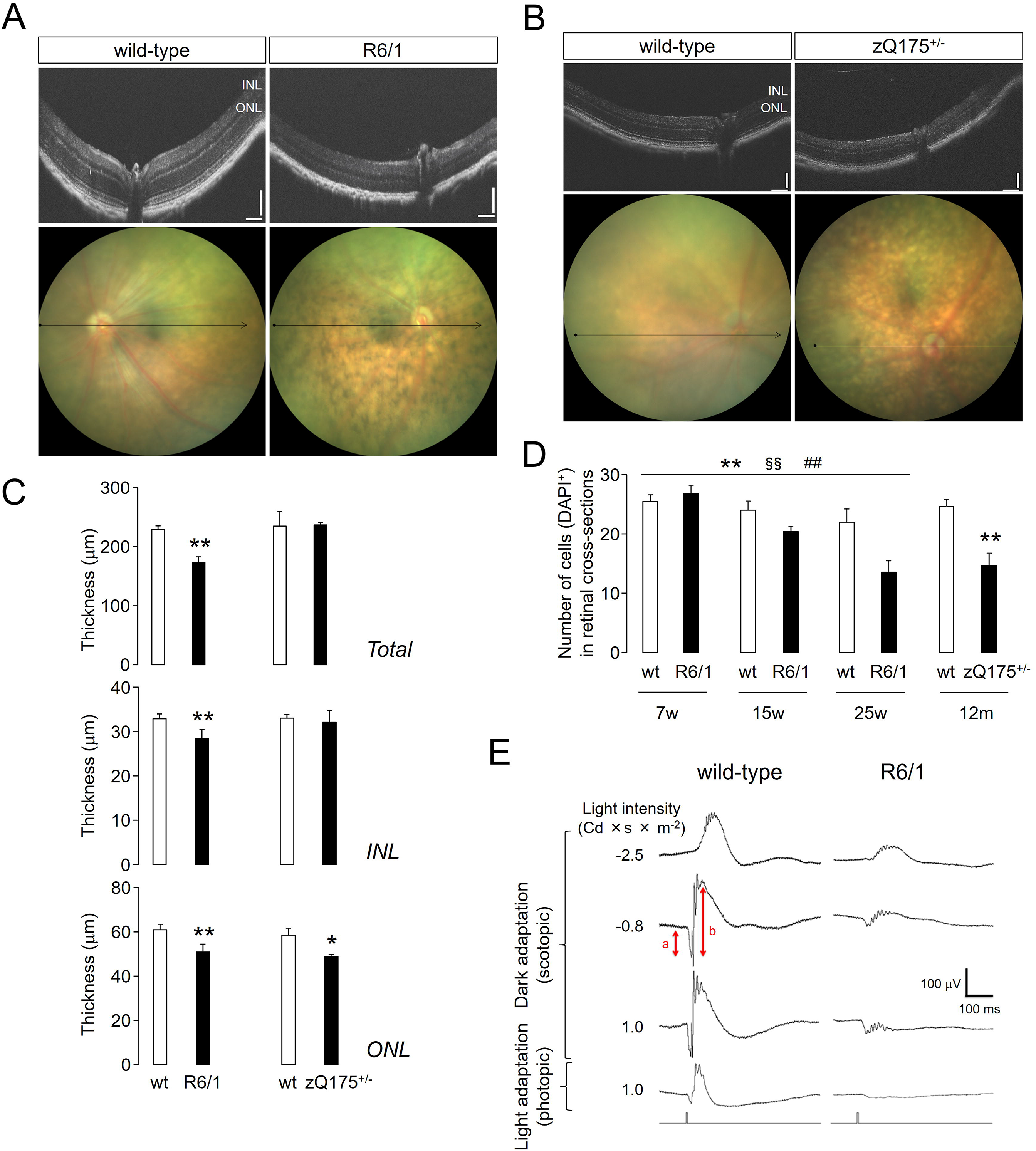
Impairment of the R6/1 and zQ175 retinas. *A-B*, Representative SD-OCT images from the fundus of a R6/1 mouse (*A*), a zQ175 mouse (*B*) and wt littermates. Arrows denote the B-scanned transects for thickness calculations. n = 2 (wt), n = 4 (R6/1), n = 3 (zQ175). Scale bars = 100 μm. *C*, Quantifications of the retinal thickness in 25-week-old R6/1 (n = 3), 12-month-old zQ175 (n = 2) and matched-age wt (n = 4 for each strain). Data are shown as mean ± SD. *, *p*<0.05; **, *p*<0.005, Student’s *t*-test between genotypes. INL, inner nuclear layer; ONL, outer nuclear layer. *D*, Quantification of DAPI-stained cells in the retina of mutant mice compared to wt across different time points, n = 3 for each genotype (7-week-old and 12-month-old animals) and n = 4 for each genotype (15 and 25-week-old animals). Data are shown as mean ± SD. **, *p*<0.005 genotype effect; §§, *p*<0.005 age effect; ##, *p*<0.005 interaction effect from ANOVA test in R6/1 and wt littermates. **, *p*<0.005, Student’s *t*-test between wild-type and zQ175 mice. *E*, Representative ERG recordings of a wild-type and R6/1 mouse, depicting a- and b-wave amplitudes. n = 4 for each genotype.

### Both the retina and striatum of R6/1 mice showed extensive transcriptional dysregulation

Next, we examined the molecular alterations underlying the impairments observed in the R6/1 retina and their relationships with the dysfunction of the most affected brain area in HD, the striatum. To this end, we screened by RNA-seq the whole mRNA expression of the retinas and striata obtained from the same mutant and wild-type mice in an earlier pathological stage (13-15 weeks-old). The differential expression analysis (adjusted *p*-value < 0.05) revealed extensive rearrangement of the R6/1 retina transcriptome related to wild-type littermates (1078 downregulated and 575 upregulated genes), resulting in more DEGs compared to the striatum of the same animals (763 downregulated and 176 upregulated genes) although comparable in magnitude and significance (Fig. 2A-B). Most of the resulting DEGs confirmed previous reports in the striatum ^27,45–47^ and in a single study conducted in the retina with an early microarray platform in the R6/2 strain ^48^, although our study increased the amount of information by 6-fold when comparing the number of DEGs.

**Figure 2.**
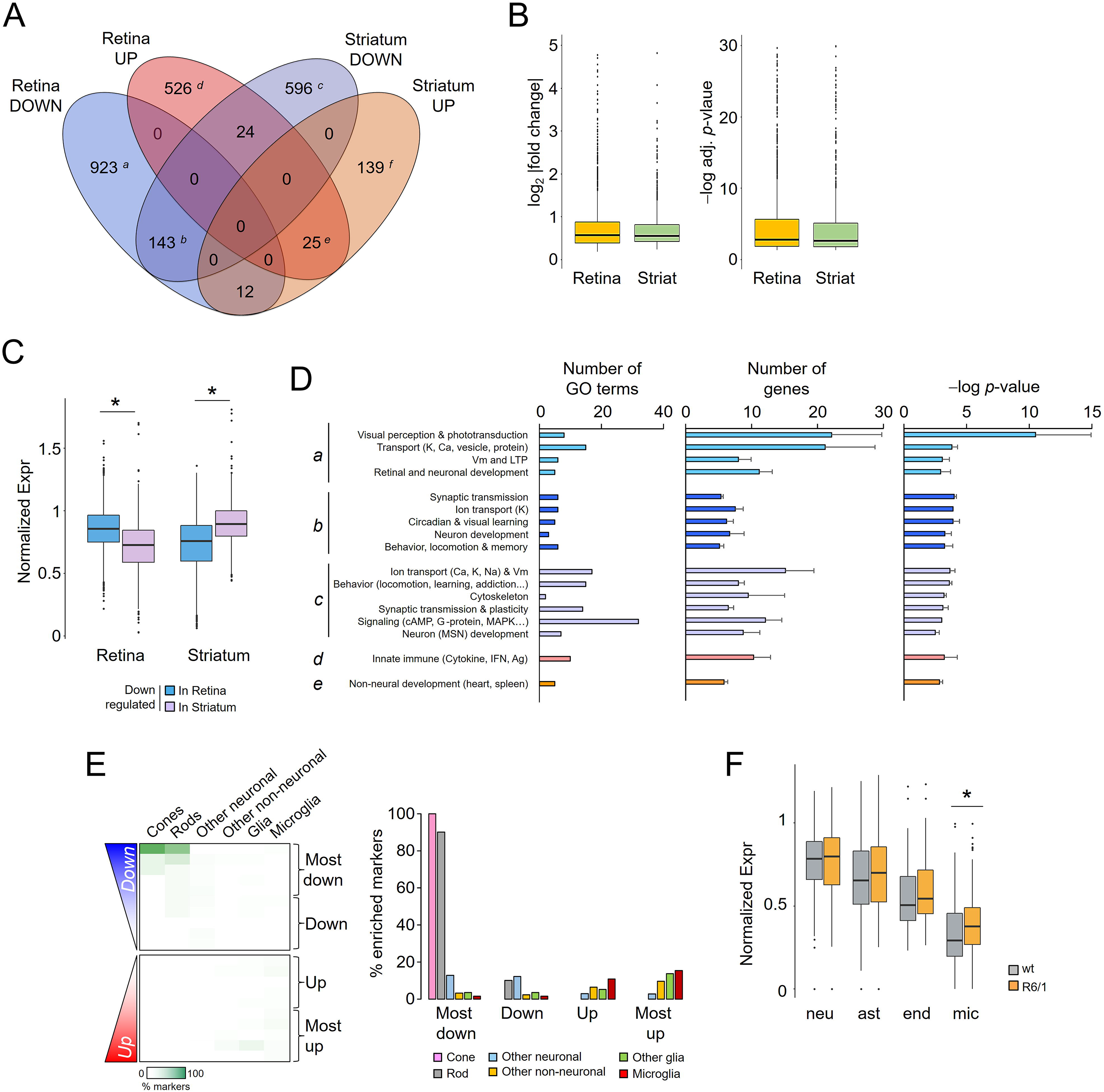
RNA-seq analysis in the retina and striatum of R6/1 mice. *A*, Venn diagram showing the number of DEGs in pairwise comparisons between wt and R6/1 mice (adj. *p*-value < 0.05). Downregulated genes were indicated as *a* (exclusive retinal), *b* (common to both tissues) and *c* (exclusive striatal), and upregulated genes as *d* (exclusive retinal), *e* (common) and *f* (exclusive striatal). In total, retinal downregulated genes consisted of subsets *a*, *b* and 12 genes; striatal downregulated genes of *b*, *c* and 24 genes; retinal upregulated genes of *d*, *e* and 24 genes; striatal upregulated genes of *e*, *f* and 12 genes. *B*, Absolute values of log_2_ fold change (left) and −log adj. *p*-value (right) in the R6/1 retina and striatum: outliers >5 and >30 were removed respectively from each panel to enable visualization of medians and quartiles of the values. *C*, Normalized basal expression of DEG in wild-type retinas and striata. *, *p*-value <0.05, Student’s *t*-test between tissues. *D*, GO enrichment analysis of DEG in the retina and the striatum of R6/1 mice. Only those terms with *p*-value < 0.001 (DAVID) are shown; not significant results were retrieved for subset *e*. Letters indicate the subsets of genes defined in *A*. Data are expressed as mean ± s.e.m. Vm, membrane potential; cAMP, cyclic AMP; MSN, medium spiny neurons; IFN, interferon; Ag, antigen. *E*, Retinal DEGs were ranked according to their significance and direction of change, and divided in bins of 100 genes. On the left, percentage of retinal cell-specific markers (see text) that were counted per bin. For “Other neuronal” we represent the average of counts from amacrine, bipolar and retinal ganglion cells, for “Other glia” the averaged counts from Müller glia and astrocytes, and for “Other non-neuronal” the averaged counts from pericytes and perivascular fibroblasts. On the right, the same data were represented as grouped percentages in a bar graph. *F*, Normalized expression of curated cell-enriched markers for neurons (neu), astrocytes (ast), endothelial cells (end) and microglia (mic) in the retina of wt and R6/1 mice. *, *p*-value <0.05, Student’s *t*-test between genotypes.

Using our significance cut-off, most of the DEGs in our RNA-seq analysis were specific to the retina and the striatum: 87.7% and 78.3%, respectively. In fact, downregulation generally affected genes that were highly expressed in the corresponding neural tissue: i.e., downregulated retinal genes were physiologically more highly expressed in the retina than in the striatum, and vice versa (Fig. 2C), in agreement with the strong tissue-enriched component of HD-associated transcriptional dysregulation (^46,49^ and references therein). This tissue specificity was also confirmed at the functional level (Fig. 2D), since we detected that downregulated genes in the R6/1 retina (subset *a*) were enriched in GO terms genes associated with visual perception, phototransduction and retinal development, whereas downregulated genes in the striatum (subset *c*) were enriched with genes linked to addiction, locomotion, and medium spiny neurons, among the expected pathways affected in HD (e.g., cAMP and G-protein-dependent). Notably, we retrieved circadian functions among the commonly altered genes in both tissues (subset *b*) that might be related to the circadian-like modulation of GABAergic interneurons and dopamine in diverse brain areas, including the retina and striatum ^50,51^. This observation should be explored in future studies considering the circadian disruption in HD patients and mice ^5,52^. In any case, neuronal functions were enriched (e.g., synaptic transmission, ion transport, neuronal development) in all subsets of downregulated genes (Fig. 2D). In contrast, upregulation was linked to distinctive phenomenologies in each tissue, as evidenced by the retrieval of nonoverlapping associated functions (i.e., innate immune response in retinal subset *d* and nonneural development in striatal subset *f*, Fig. 2D) and different putative regulatory mechanisms (Supplementary Fig. S2A, B). The upregulation of immune-related genes in the retina was consistent with the fundus anomalies observed in the R6/1 mice (Fig. 1A-B).

To explore in further detail the retinal transcriptional profile of the R6/1 strain, we interrogated the cell types that were the most affected according to our differential expression analysis. To this end, we used the markers obtained from a scRNA-seq analysis performed in the mouse retina that were assigned to specific neuronal and nonneuronal cells ^30^ (see Materials and Methods for further details). The presence of these markers among the DEGs inferred the main affectation of photoreceptor cells as specific genes of cones (e.g., *Opn1sw*, *Arr3*) and rods (e.g., *Rho*, *Gnat1*) were the most downregulated genes in R6/1 retinas compared to wild-type littermates, followed by the upregulation of microglia (e.g, *Trf*, *H2-K1*, *Ctss*) and astrocytic markers (e.g., *A2m*, *Gfap*) (Fig. 2E), in agreement with the GO enrichment result in Fig. 2D. These affectations were not sufficient to alter the levels of general neuronal and astrocytic markers at this age of sampling; in contrast, we observed a significant microglial activation (Fig. 2E).

### R6/1 retinas showed glial activation

Our RNA-seq analysis suggested that local inflammatory processes took in the retina, but not in the striatum, of symptomatic R6/1 mice, in agreement with a TFBS prediction in which DNA motifs for inflammatory transcription factors (i.e., IRFs and STATs) ^53^ were highly enriched in the genes that were upregulated in the R6/1 retina (Supplementary Fig. S2). This retina-specific neuroinflammation was not due to either a higher CAG instability of the transgene (Fig. 3A) or higher expression of the R6/1 transgene in the mutant retina related to the striatum (Fig. 3B), which might accelerate the progression of the pathology in the former tissue.

**Figure 3.**
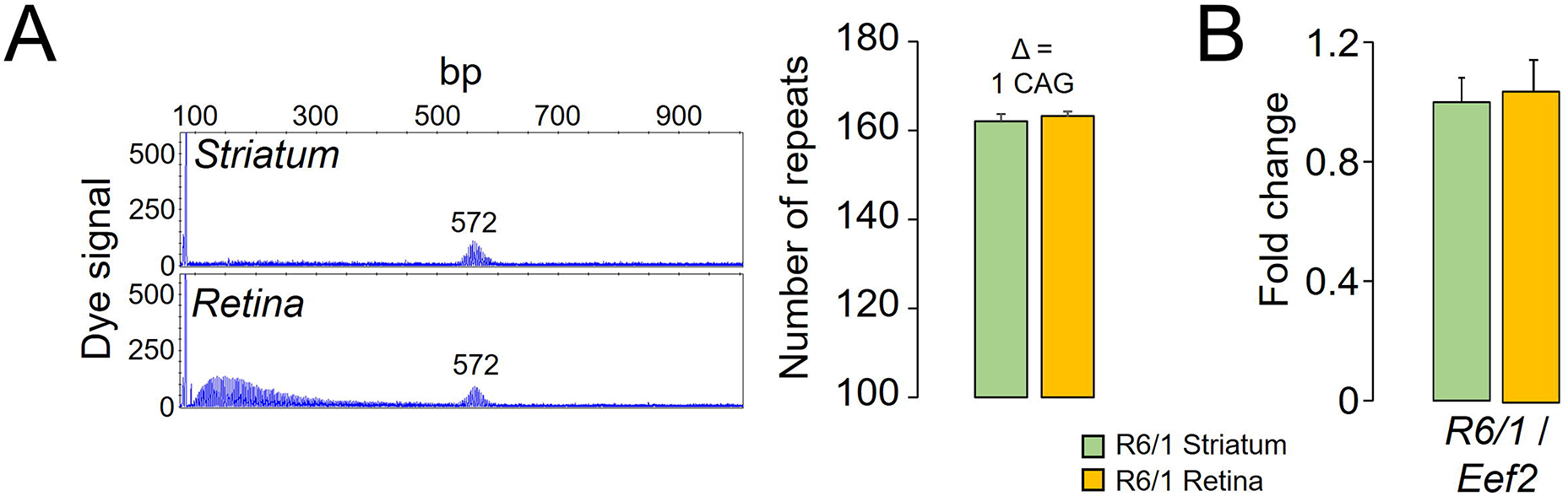
Number of CAG repeats and R6/1 transgene expression in the R6/1 retina and striatum. *A*, Representative electropherograms of DNA fragment analysis in the striatum and the retina of a mutant mouse (left panel). Quantification of the results from ten R6/1 mice (right panel): only a net single CAG repeat was different between both tissues (Δ). *B*, RT-qPCR assays of R6/1 retinas and striata (n =13 each) from 5 and 7-week-old mice, prior to any potential cell loss that might influence the result. Transgene expression was normalized by *Eef2* values as this gene was equally expressed in both tissues in the RNA-seq data (Supplementary Fig. S3A). Data are expressed as mean ± s.e.m.

Since retinal markers used in Fig. 2D was conducted in the basal state, this analysis did not differentiate the activation states of resident glial cells. To determine whether the retinal neuroinflammation was associated with glial activation, we used external datasets to obtain the most significant genes (top 250) in the pairwise comparison between isolated astrocytes (Aldh1l1^+^) from control and injured mice (combining LPS injection and middle cerebral artery occlusion ^33^): whereas the upregulation signature was linked to astrocytic activation and gliosis, downregulated genes indicated overexpression in resting astrocytes. In a similar manner, we also obtained the top 250 genes from isolated microglia (CD11b^+^-CD45^+^) at different stages of neurodegeneration triggered by p25 induction; thus, we retrieved the signatures for homeostatic/basal, early and late activated microglia under ongoing neurodegenerative processes ^36^. We identified examples of all sets of genes related to basal and activated glia to be significantly upregulated in the R6/1 retina compared to wild-type littermates, suggesting the coexistence of different stages of glial activation (Fig. 4A, top panels). In contrast, the R6/1 striatum did not show any increase in glial markers (Fig. 4A, bottom panels). To gain further insights regarding the type of glial activation, we examined the changes occurring in markers ascribed to two recently described subtypes of reactive astrocytes: neurotoxic A1 and neuroprotective A2 ^37^. We found in the retina of R6/1 mice the activation of 25% of A1-astrocytic and 38.5% pan-astrocytic markers, indicating that the detected gliosis was associated with an inflammatory deleterious response (Fig. 4B). In the case of microglia, we were unable to distinguish the two classical polarization states M1 (proinflammatory) and M2 (antiinflammatory) using markers described elsewhere ^54^.

**Figure 4.**
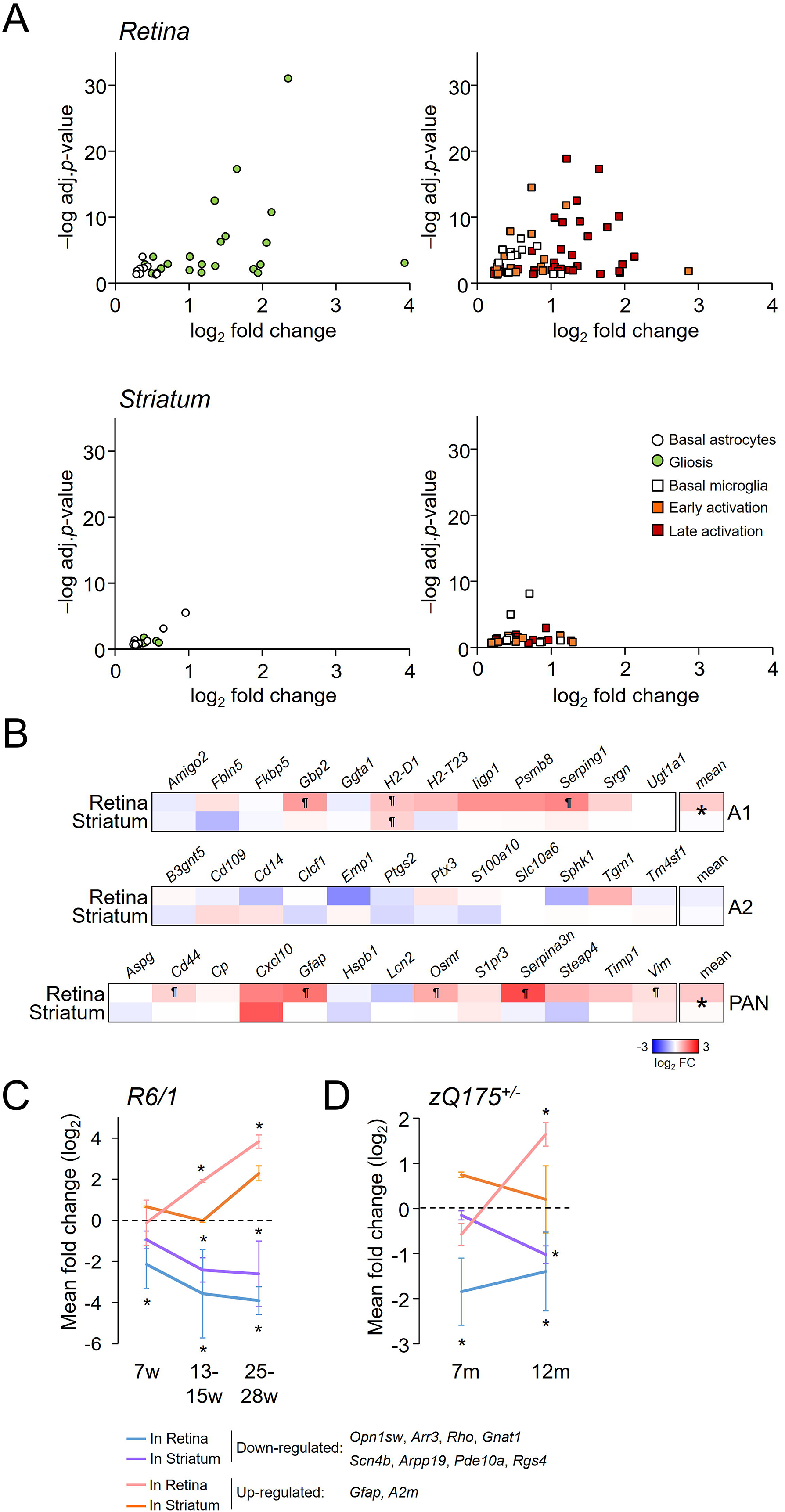
Presence of a glial signature in the R6/1 and zQ175 retinas. *A*, Plots showing the upregulated genes in R6/1 retina (top panels) and striatum (bottom panels) according to magnitude (log_2_ fold change) and significance (−log adj. *p*-value) of change that are markers for astrocytes (left panels) and microglia (right panels) at different stages of activation (see text for further details). For striatal genes no significance cut-off was applied to permit the analysis of the same number of genes as in retina to facilitate the comparison between both tissues. *B*, Heatmap plot of the fold changes of DEGs belonging to the A1, A2 and panastrocytic signatures (see text). ¶, adjusted *p*-value < 0.05 from the RNA-seq analysis. *, *p*-value < 0.05, Mann Whitney *U*-test between the fold changes means of both tissues. *C-D*, Summary of the RT-qPCR results represented in Supplementary Fig. S4. Fold changes of selected down- and up-regulated genes in R6/1 (*C*) and zQ175 (*D*) were averaged (mean ± SD) in each sampling time point. Dash line denotes the wt reference. *, *p*-value < 0.05, Student’s *t*-test between genotypes for all down- or up-regulated genes.

Retinal neuroinflammation was apparently restricted to glial cells since additional inflammatory markers were not induced in our RNA-seq data. For example, most adaptive immune markers for T- and B cells were not detected in the retinal transcriptomes or were expressed at low levels, independently of the genotype (Supplementary Fig. S3A). In addition, other genes associated with inflammation were also lowly expressed (such as the classical proinflammatory cytokines *Il1b*, *Il6* and *Tnf*) or did not show differences between genotypes (such as the inducible heme oxygenase by mitochondrial dysfunction *Hmox1*), as confirmed by RT-qPCR assays in independent samples (Supplementary Fig. S3B).

To establish the time-frame in which R6/1 retinas exhibited evident glial activation, we analysed the changes in the gene expression of two of the most significantly upregulated astrocytic genes, *A2m* and *Gfap*, at different stages of the pathology: prodromal (7 weeks old), symptomatic (13-15 weeks old) and advance symptomatic (25-28 weeks old). Upregulation of these markers was significant after the downregulation of neuronal-related genes in the R6/1 retina; nonetheless, all examined genes were significantly altered in the intermediate age of sampling (Fig. 4C and Supplementary S4A, B). In contrast, we only observed a nonsignificant upregulation trend for these two astrocytic genes in the R6/1 striatum compared to wild-type striatum at advanced stages of the disease (Fig. 4C and Supplementary S4C, D). Independent samples from 24-week-old animals confirmed the lack of significance for the upregulation of *Gfap* and *A2m* in the R6/1 striatum (Supplementary Fig. S4E). A similar behaviour was observed in the zQ175 mice, in which changes in 12-month-old mice were comparable to those occurring in R6/1 mice 13-15 weeks old (Fig. 4D and Supplementary S3F, G). Notably, the CAG expansion in homozygosis only accelerated the changes of striatal genes at an earlier time point in the KI model (7 months old, Fig. Supplementary S3F, G).

Because these analyses were performed using bulk homogenates, we investigated the expression of GFAP by immunofluorescence to obtain spatial sensitivity of astrocytic activation. Reactive GFAP was detected in retinal sections from R6/1 mice beginning at 7 weeks of age with increased labelling in the following weeks, first in the feet of the ganglion cell layer (GCL), extending to the outer plexiform layer (OPL) and later continuing into the inner nuclear (INL) and inner plexiform layers (IPL) in R6/1 mice at 25 weeks of age (Fig. 5A). Reactive astrogliosis was corroborated in the 12-month-old zQ175 retinas (Fig. 5B). To analyse the microglial cell expression and activation, we determined the time-course staining pattern of the specific reactive microglial marker Iba-1. Immunopositive Iba-1 cells were detected mainly in both plexiform retinal layers of both HD mouse models (Fig. 5C-D). We observed a nonsignificant elevation in the total number of microglial cells during pathology progression but, more interestingly, there was a prominent shift towards an amoeboid-activated phenotype detriment to ramified-resting morphology in the retinas of 25-week-old R6/1 mice, indicating ongoing microglial activation (Fig. 5C) that explained the elevated levels of general microglial markers documented in Fig. 2E. This shift was also observed in the 12-month-old zQ175 retina (Fig. 5D). The localization of activated microglial cells identified the OPL as the most damaged layer, potentially explaining the loss of synaptic connection between the ONL and the INL determined by ERG recordings and the lack of TUNEL-positive cells since retinal apoptotic cells might be removed by the increased phagocytic capacities of activated microglia ^55^.

**Figure 5.**
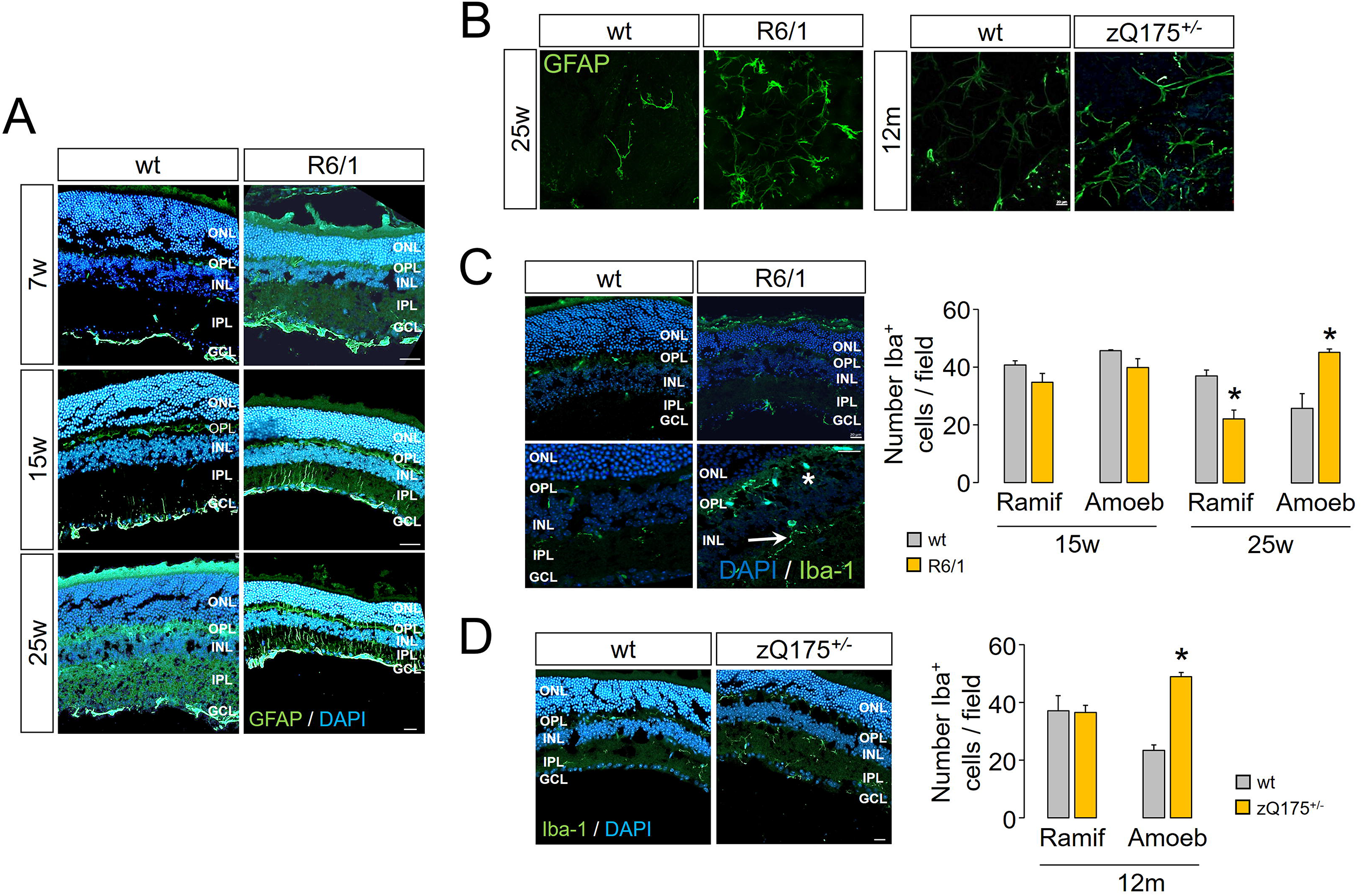
Glial activation in the R6/1 and zQ175 retinas. *A*, Representative immunofluorescence stainings across the indicated time points of the gliosis marker GFAP in R6/1 and wt littermates. *B*, Representative whole mounts of R6/1, zQ175 and wt retinas with GFAP staining. *C-D*, Representative immunofluorescence staining of the microglia marker Iba-1 for R6/1 (*C*) and zQ175 (*D*). Right panels, quantification of the Iba-1^+^ cells distinguishing their morphology in ramified and amoeboid in R6/1 (*C*) and zQ175 (*D*) retinas compared to wt littermates. N = 2 per genotype, age and model; n = 3 – 5 slices per animal, n = 3 fields per slice; field area = 84100 μm^2^. Data are expressed as mean ± SD.*, *p*-value < 0.05, Student’s *t*-test between Iba-1^+^ cell subtypes. Green, glial marker; blue, DAPI staining. Scale = 20 μm. ONL, outer nuclear layer; OPL, outer plexiform layer; INL, inner nuclear layer; IPL, inner plexiform layer; GCL, ganglion cell layer; Ramif, ramified; Amoeb, amoeboid.

### The autophagy processes were differentially modulated in the retina and striatum of R6/1 mice

On the basis that autophagy is known to be altered in HD ^56^ and is involved in inflammation resolution due to its clearance role ^57,58^, we compared the dysfunctions in autophagy processes between the retina and the striatum of R6/1 mice and their wild-type littermates. After inspection of the RNA-seq results, we observed the significant deregulation of several components of the autophagic system in nearly all the categories summarized in ^38^ in both the retina and the striatum of R6/1 mice (Fig. 6A). The most notable difference was observed between retinal down- and upregulated genes in the lysosomal category (χ^2^= 12.13, d.f. = 5, *p*-value = 0.033, Fig. 6A). This category included genes involved in lysosomal biogenesis that were in general upregulated in the retina compared to the striatum of the same mice (Fig. 6B).

**Figure 6.**
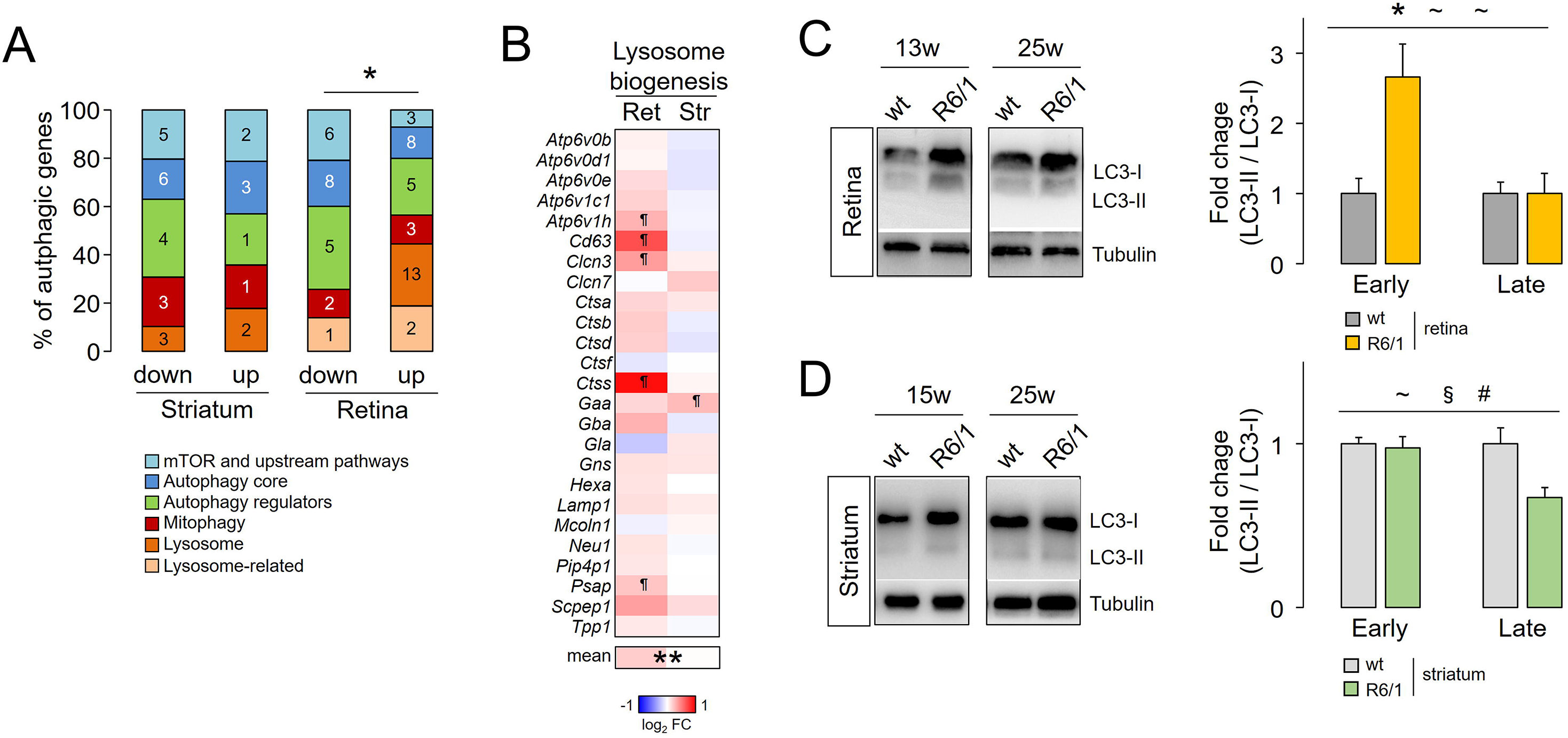
Autophagy modulation in the retina and the striatum of R6/1 mice. *A*, Proportion of autophagy-related genes belonging to different categories (see text), considering the number of genes within each category and within DEGs in R6/1 tissues. Numbers indicate the number of significant DEGs obtained in the RNA-seq analysis. *B*, Heatmap plot of the fold changes of DEGs belonging to the “Lysosome biogenesis” signature described in the text. ¶, adjusted *p*-value < 0.05 from the RNA-seq analysis. **, *p*-value < 0.005, Mann Whitney *U*-test between the fold changes means of both tissues. *C-D*, Protein extracts from the retina (*C*) and striatum (*D*) of R6/1 and wild-type littermates were analyzed by western blotting assays with antibodies against LC3II/I and α-tubulin as loading control. Retina early, 7/9/15-week-old, n = 4 (wt) and n = 6 (R6/1); Retina late, 21/25-week-old, n = 3 (wt) and n = 4 (R6/1); striatum early, 5/13-week-old, n = 6 (wt) and n = 9 (R6/1); striatum late, 25-week-old, n = 4 (wt) and n = 5 (R6/1). Data are expressed as mean ± s.e.m. *, *p*<0.05 genotype effect; §, *p*<0.05 age effect; #, *p*<0.05 interaction; ~, *p*<0.1 in any effect, ANOVA test in R6/1 and wt littermates. More blots are shown in Supplementary Fig. S6.

We next analysed by western blotting assays the conversion of the cytosolic form of LC3 (LC3-I) into the LC3-phosphatidylethanolamine conjugate (LC3-II), which is recruited to autophagosomal membranes, as a marker of autophagy activation. Based on their similarity, we grouped the results for R6/1 at early (5-15 weeks old) and advance stages (21-25 weeks old). We detected a significant increase in the LC3-II / LC3-I ratio in the retinas from early R6/1 mice compared with age-matched wt mice in response to the mHTT insult, but the LC3-II / LC3-I ratio was later similar in advanced R6/1 mice (Fig. 6C), indicating that autophagic influx was delayed ^59^. The inability to activate the autophagic influx in the mutant striatum was already observed in early stages, leading to the accumulation of LC3-I in advanced stages (Figure 6D) in agreement with other reports ^60,61^. Overall, these results indicated that autophagy activity was differentially impaired in the retina and striatum of R6/1 animals.

## Discussion

The present work defines for the first time the relationship between the molecular alterations in the retina and striatum in HD mouse models. Retinopathy in HD has been described as a late event in the pathology, based on the normal fundus and minimal ERG anomalies in young R6/1 animals ^17,20^. However, degeneration of specific cell types and altered retinal responses to light can occur in the R6/2 model prior to the onset of motor impairment and body weight loss ^18,19^. Another polyQ disorder caused by an aberrant expansion of CAG triplets in the *ATXN7* gene, spinocerebellar ataxia 7 (SCA7), is characterized by retinal degeneration and dystrophy, leading to visual anomalies preceding motor symptoms ^62^. HD and SCA7 animal models display common features of retinal anomalies at the level of morphology, light response and gene expression ^20,48,63^, retinal affectation may share similar progression in both disorders and might manifest in early symptomatology.

Although gliosis has been previously reported in the R6/1 retina ^17^, we extended this observation to heterozygous zQ175 KI mice, which better resemble the pathology in humans since the disease progression is much slower and comprises the loss-of-function component ^23^. We also provided the first evidence of microglial activation in the retina of two different HD models as a marker of neuroinflammation. Microglia comprise the resident phagocyte population in the CNS that can have both protective and deleterious effects, with anti-inflammatory and repair responses during the early phase of neurodegeneration but with harmful effects on neurons in prolonged activation in chronic inflammation ^64–67^. In our RNA-seq analysis, we were able to detect different states of microglial activation, consistent with the hyperreactivity reported in microglia derived from pluripotent stem cells and blood-isolated myeloid cells of HD patients ^68,69^. Reactive microglia induce the proinflammatory A1 astrocyte subtype, which is found in patients’ brains affected by different neurodegenerative disorders including HD ^37^. In the R6/1 retina, we observed the induction of some of these astrocytic markers; however, our data also suggested that astrocytic diversity falls beyond the dichotomous classification in A1 and A2 as reported in patients ^70^.

Our striatal results were in agreement with the absence of overt neuroinflammation in HD models ^27,71^. However, in HD patients activated glia and induction of inflammatory markers are well detected in cortical and basal ganglia regions ^72–74^. This discrepancy is exemplified in a recent study that compared the gene expression profiles of murine R6/1 and zQ175 striata with those of human caudate nuclei from HD patients at a single nucleus resolution ^75^. The induction of pan- and A1-astrocytic markers in human HD samples was reminiscent of our own results in the retina of R6/1 mice, suggesting that mHTT-expressing retinas might better reproduce the conditions of HD-associated neuroinflammatory processes. Since glia show regional-dependent heterogeneity at the basal and activated levels ^36,76,77^, it could be of interest to further compare their intrinsic properties in both tissues.

Tightly connected to inflammatory processes, autophagy delivers aberrant organelles and macromolecules in double-membrane vesicles to lysosomes for degradation and recycling ^59^. This process alleviates the cellular damage induced by neuroinflammation ^78^, and modulates the functionality of immune and glial cells ^79^. In HD autophagy becomes severely disrupted at several steps (e.g., cargo recognition, autophagosome formation, maturation and fusion to lysosomes), not only due to the toxic effects of the polyQ peptide but also because of the reduced regulatory activity of physiological HTT over autophagy, affecting the clearance of mHTT and contributing to its accumulation in cells ^56^. We confirmed that autophagy was altered in the retina and striatum of R6/1 mice, but apparently showed different rates of autophagosome accumulation, in which autophagic influx was stopped earlier in the R6/1 striatum than in the retina. The latter was transiently able to respond to mHTT. Whether the upregulation of lysosomal components (including the microglial cathepsin *Ctss*) is part of a compensatory mechanism for the HD-associated deficiency in autophagy deserves further exploration. Based on our results, we propose that the retinas of HD mouse models can serve as a model to analyse HD-linked neuroinflammation, allowing for the use of *ex vivo* retina cultures from HD mouse models to elucidate mechanisms of neurodegeneration and to evaluate potential therapeutic approaches ^80^, as research models that are evolutionarily closer than HD *Drosophila* ommatidia ^81–84^.

In conclusion, the retinas of HD mouse models show profound morphological and functional abnormalities that are accompanied by a dramatic transcriptional dysregulation and glial activation, suggesting that retinal affectation may be of higher relevance in HD than envisaged. Studying this activation can provide novel insights regarding the role of astrocytes and microglia during HD neuroinflammation.

## Abbreviations

*A2m*: Alpha-2-macroglobulin
*Actb*: Actin beta
*Arr3*: arrestin 3
cAMP: cyclic adenosine monophosphate CNS, central nervous system
*Ctss*: Cathepsin S
DAPI: 4’-6-diamidino-2-phenylindole DEG, differentially expressed gene ERG, electroretinogram
FDR: false discovery rate
*Gapdh*: Glyceraldehyde-3-phosphate dehydrogenase GCL, ganglion cell layer
GFAP: glial fibrillary acidic protein
*Gnat1*: G-Protein subunit alpha transducin 1
*H2-K1*: H-2 class I histocompatibility antigen, K-B alpha chain HD, Huntington’s disease
*Hmox1*: Heme oxygenase 1 HTT, huntingtin
Iba-1: ionized calcium binding adaptor molecule 1
*Il1b*: interleukin 1b
*Il6*: interleukin 6
INL: inner nuclear layer IPL, inner plexiform layer
IRF: interferon regulator factor KI, knock-in
ONL: outer nuclear layer OPL, outer plexiform layer
*Opn1sw*: Opsin 1, short wave sensitive
*Rho*: rhodopsin
RNFL: retinal nerve fibre layer
RT-qPCR: retrotranscription and quantitative PCR scRNA-seq, single cell RNA sequencing
SD-OCT: spectral domain optical coherence tomography STAT, Signal transducer and activator of transcription
*Tbp*: TATA-box binding protein
TFBS: transcription factor binding site
*Tnf*: Tumor necrosis factor
*Trf*: Transferrin
*TUNEL*: terminal deoxynucleotidyl transferase-mediated dUTP nick-end labelling UHDRS, Unified Huntington’s Disease Rating Scale
wt: wild-type

## Acknowledgements

We thank Rocío Pérez-González for critical reading of the manuscript and José J. Lucas for providing mice to establish our zQ175 colony.

## Authors’ contribution

Conceptualization, Funding acquisition, Project administration, Resources and Supervision: Luis M. Valor and Ana I. Arroba. Data curation, Formal analysis and Software: Luis M. Valor and Fátima Cano-Cano. Investigation: Andrea Gallardo-Orihuela, Fátima Cano-Cano, Francisco Martín-Loro, Maria del Carmen González-Montelongo, Irati Hervás-Corpión, Pedro de la Villa, Lucía Ramón-Marco, Laura Gómez-Jaramillo, Ana I. Arroba and Luis M. Valor. Methodology: Ana I. Arroba, Luis M. Valor and Pedro de la Villa. Software: Fátima Cano-Cano and Luis M. Valor. Validation: Francisco Martín-Loro, Maria del Carmen González-Montelongo and Lucía Ramón-Marco. Visualization: Luis M. Valor, Francisco Martín-Loro and Ana I. Arroba. Writing - original draft: Luis M. Valor. Writing - review & editing: Ana I. Arroba, Maria del Carmen González-Montelongo, Fátima Cano-Cano, Francisco Martín-Loro, Irati Hervás-Corpión, Andrea Gallardo-Orihuela, Laura Gómez-Jaramillo and Luis M. Valor.

## Data statement

The RNA-seq data can be downloaded from the Gene Expression Omnibus (GEO) database using the accession number GSE216520.

