## Supplemental Figures for "Retinal affectation in Huntington’s disease mouse models concurs with a local innate immune response"

**Supplementary Figure S1. mHTT and TUNEL analysis in HD retinas.** *A*, Representative immunofluorescence images of 25-week-old wt and R6/1 retinas against mHTT (green) and counterstained with DAPI (blue). Scale = 20 μm (inset, 5 μm). *B*, Representative TUNEL images in matched-age wt, R6/1 and zQ175 (n = 3 each genotype). Only a couple of spare cells were observed to be positive for DNA fragmentation. Rd10 retinas were used as positive control. Scale = 20 μm.

**Supplementary Figure S2. Prediction analysis of DNA motifs at the promoters of R6/1 deregulated genes.** *A*, Pscan enrichment of Jaspar-based TFBS at the -950/+50 promoter regions in the subsets of DEG as defined in Figure 2A. TFBS in all subsets were ranked according to the Z-scores obtained for exclusive retinal DEG (*a* and *d* subsets). Inset denotes the Spearman correlation coefficients (ρ) between subsets of DEG. *B*, Sum of the numbers of significant TFBS ($p < 0.05$, Pscan) that were specific or common (overlapping) to any subset of DEG, after comparing *a*, *b*, *c* (down) or *d*, *e*, *f* (up). Note that the differential result between down- and up-regulated genes was not influenced by the numbers of TFBS as they were similar between both groups of DEG. *C*, Z-scores from the enrichment analysis of TFBS related to inflammation (IRF, STAT) in the subsets of DEG. *, *p*-value < 0.05; **, *p*-value < 0.005 (Pscan).

**Supplementary Figure S3. Gene expression analysis of additional inflammatory markers.** *A*, Table showing the normalized counts of all samples (baseMeans of Deseq2

results) for specific markers for B-, T, and NK-cells. The housekeeping *Eef2* is indicated as an example of a well expressed gene. NA, not available due to very low expression. *B*, Unnormalized $C_T$ values of the qPCR for *Il1b*, *Il6*, *Tnf* and *Hmox1* in the retina and the striatum of R6/1 and wild-type littermates, using the same threshold; only *Hmox1* was well expressed beyond noise, and fold change between genotypes was calculated. Data are expressed as mean ± s.e.m.

**Supplementary Figure S4. RT-qPCR assays in independent samples of HD mice and control littermates.** *A-B*, Time-course analysis of the downregulation (*A*) and upregulation (*B*) of selected genes in the R6/1 retina compared to wild-type littermates. 7 weeks, n = 6 (wt) and n = 8 (R6/1); 13-15 weeks, n = 5 (wt) and n = 7 (R6/1); 25-28 weeks, n = 3 per genotype. *C-D*, The same analysis in the R6/1 and wild-type striata. 7 weeks, n = 7 (wt) and n = 8 (R6/1); 13-15 weeks, n = 7 (wt) and n = 9 (R6/1); 25-28 weeks, n = 3 per genotype. *E*, RT-qPCR results for *Gfap* and *A2m* in independent 24-week-old samples: n = 6 (wt), n = 5 (R6/1). *F-G*, The same analysis in the zQ175 mice and wild-type littermates. At the early time point (7 months) we included homozygous mice for the CAG expansion to check whether there was a possible exacerbation of the transcriptional dysregulation. 7 months in each tissue, n = 5 (wt), n = 4 ($zQ175^{+/-}$) and n = 4 ($zQ175^{+/+}$); 12 months in striatum, n = 6 per genotype; in retina, n = 5 (wt) and n = 7 ($zQ175^{+/-}$). Data are expressed as mean ± s.e.m. * $p<0.05$, ** $p<0.005$, genotype effect; § $p<0.05$, §§ $p<0.005$, age effect; # $p<0.05$, ## $p<0.005$, interaction effect; ~, $p<0.1$ in any effect, ANOVA test.

**Supplementary Figure S5. Raw western blots of Figure 6.** *A*, retina, *B*, striatum.

Supplementary Figure S1. mHTT and TUNEL analysis in HD retinas

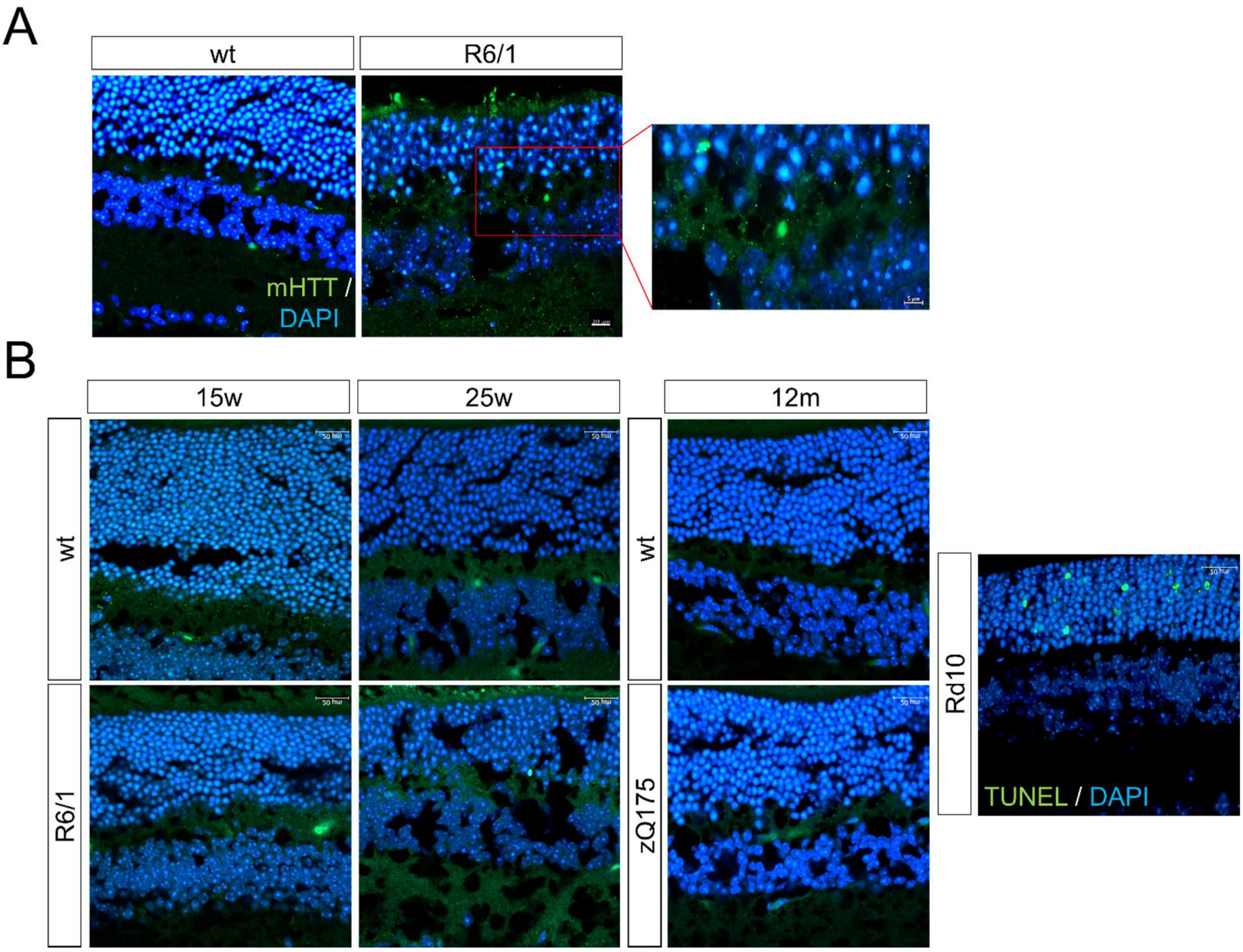

Supplementary Figure S2. Prediction analysis of DNA motifs at the promoters of R6/1 deregulated genes

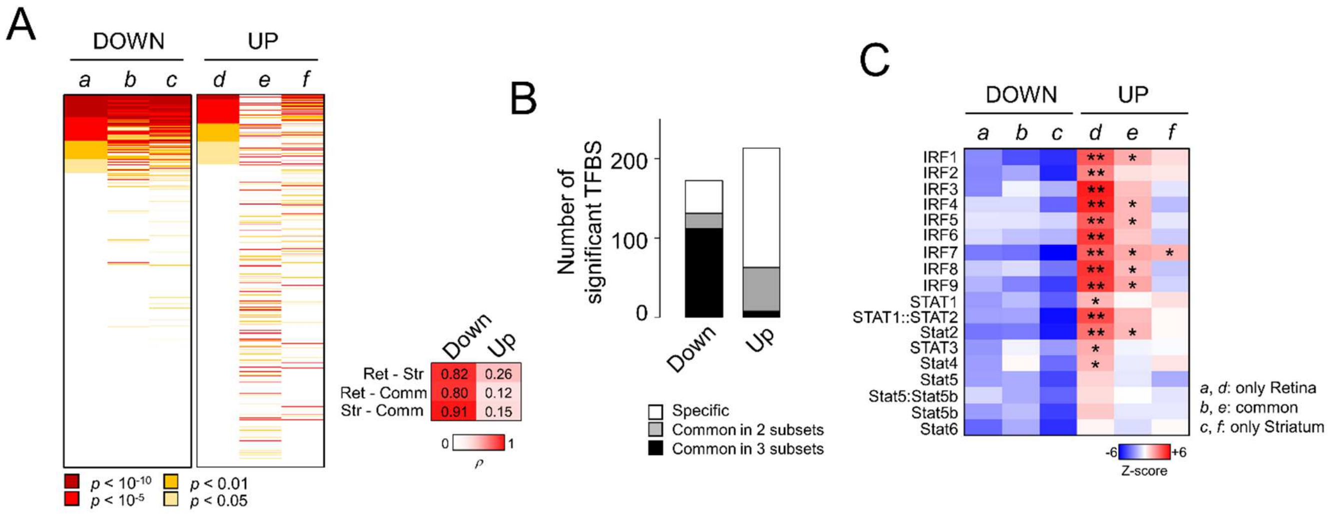

Supplementary Figure S3. Gene expression analysis of additional inflammatory markers

A

| Gene | Marker | Retina | Striatum |
|---|---|---|---|
| *Cd22* | B-cell | NA | 6.7 |
| *Cd80* | B-cell | NA | 7.5 |
| *Cd86* | B-cell | 4.6 | 46.0 |
| *Cd180* | B-cell | 10.5 | 83.8 |
| *Cr2* | B-cell | 5.3 | 10.6 |
| *Fcer2a*/CD23 | B-cell | NA | NA |
| *Ms4a1*/CD20 | B-cell | NA | 2.3 |
| *Tnfsf4*/CD252 | B-cell | NA | NA |
| *Cd3d* | NK & T-cells | NA | NA |
| *Cd3e* | NK & T-cells | NA | NA |
| *Cd3g* | NK & T-cells | NA | NA |
| *Cd247* | NK & T-cells | 2.9 | 2.3 |
| *Btla* | T-cell (CD4+) | 12.5 | 18.9 |
| *Cd4* | T-cell (CD4+) | 1.9 | 550.9 |
| *Cd40lg* | T-cell (CD4+) | NA | NA |
| *Cd69* | T-cell (CD4+) | NA | NA |
| *Cd7* | T-cell (CD4+) | NA | NA |
| *Ctla4* | T-cell (CD4+) | NA | 2.0 |
| *Icos* | T-cell (CD4+) | NA | NA |
| *Il2ra*/CD25 | T-cell (CD4+) | NA | 6.6 |
| *Il7r*/CD127 | T-cell (CD4+) | NA | 9.7 |
| *Pdcd1* | T-cell (CD4+) | NA | NA |
| *Sell*/CD62l | T-cell (CD4+) | 2.1 | 3.3 |
| *Tnfrsf9*/CD137 | T-cell (CD4+) | 7.7 | 15.1 |
| *Tnfrsf4*/CD134 | T-cell (CD4+) | 8.3 | 8.9 |
| *Cd5* | T-cell (CD8+) | NA | NA |
| *Cd8b1* | T-cell (CD8+) | NA | NA |
| *Cd8b2* | T-cell (CD8+) | NA | NA |
| *Cd27* | T-cell (CD8+) | NA | NA |
| *Cd28* | T-cell (CD8+) | 2.5 | 4.3 |
| *Eef2* | Housekeep. | 17345.4 | 17449.8 |

B

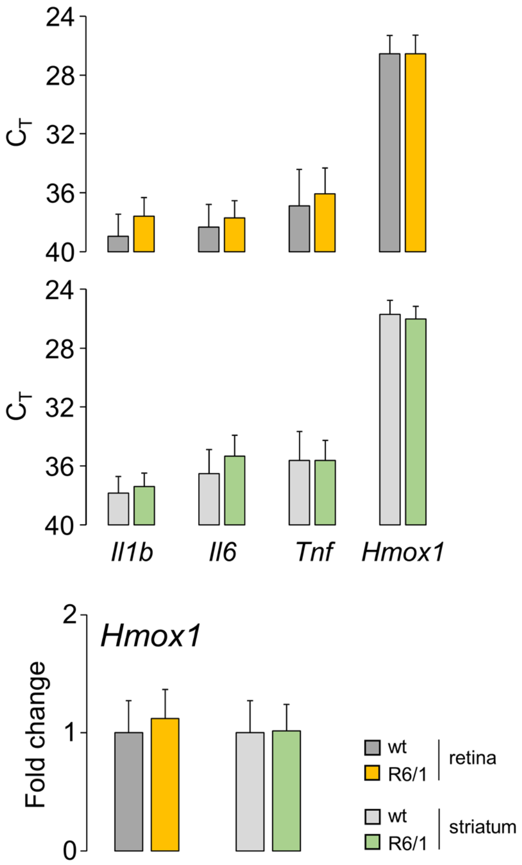

Supplementary Figure S4. RT-qPCR assays in independent samples of HD mice and control littermates

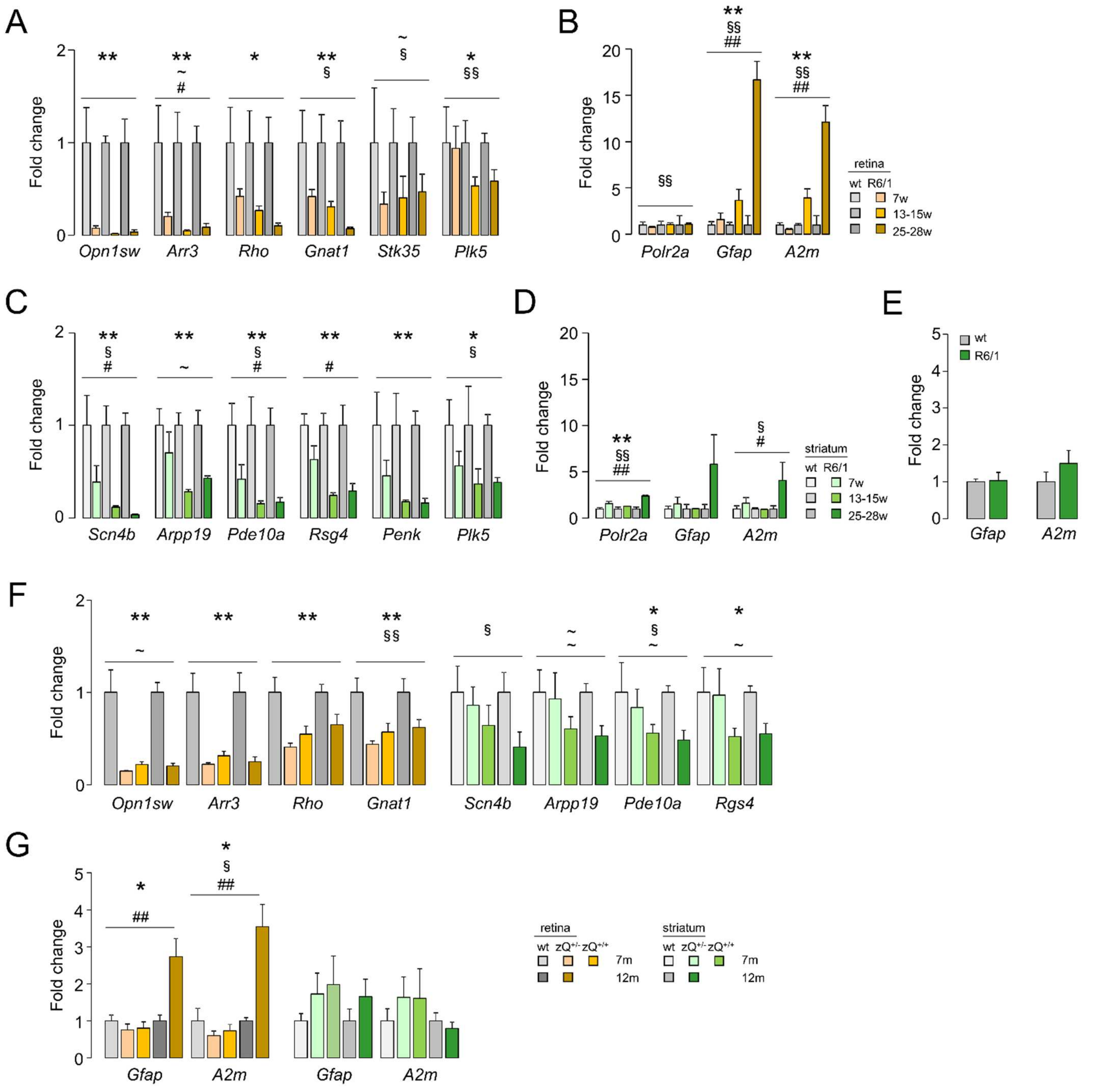

Supplementary Figure S5. Raw western blots of Figure 6

A

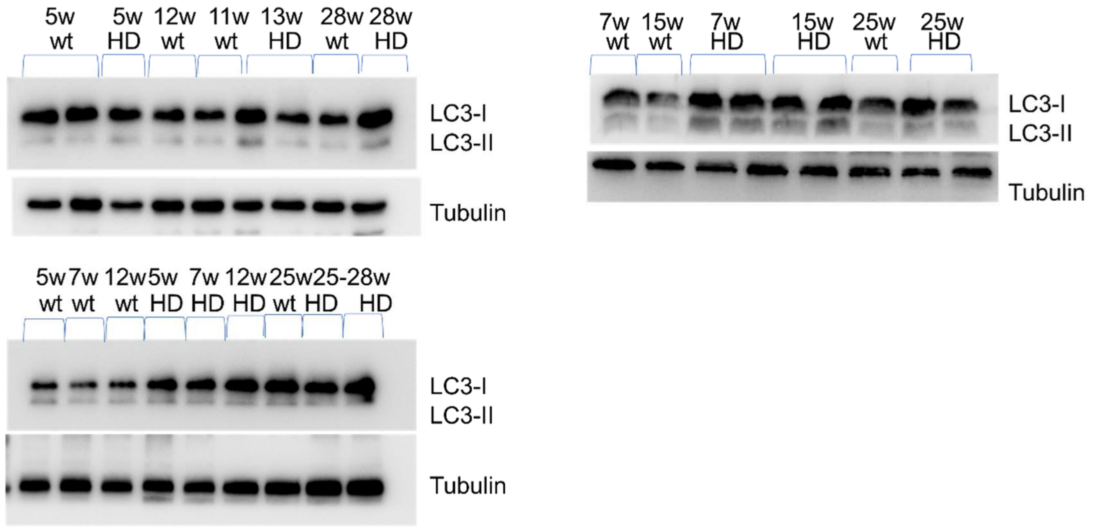


B

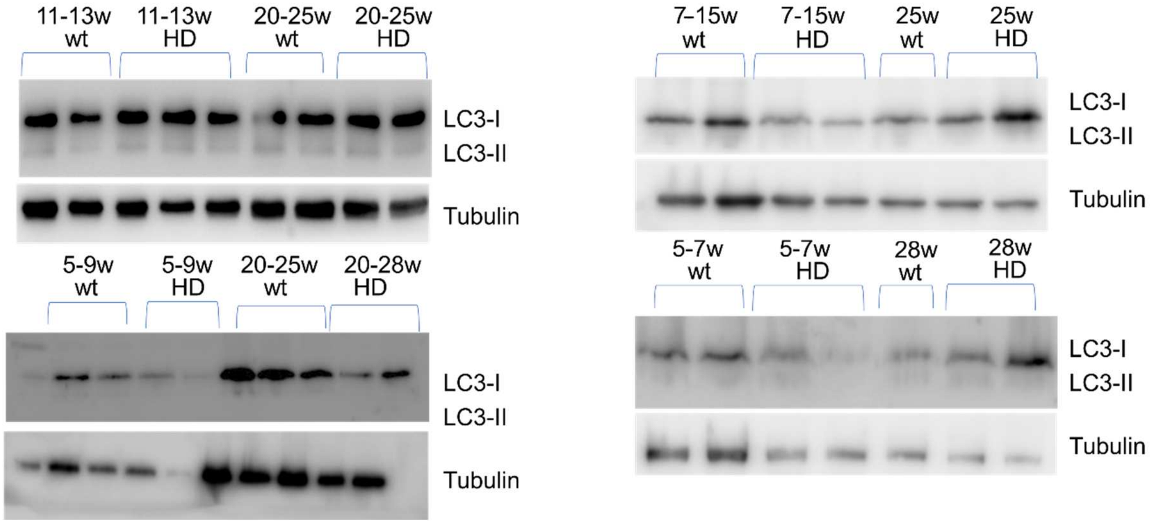
